# Intrathymic progenitor cell transplantation restores T cell development through ILC3-TEC crosstalk

**DOI:** 10.1101/2025.11.10.687563

**Authors:** Alice Machado, Chloé Houques, Marie Pouzolles, Elisa Evain, Valérie Dardalhon, Christelle Harly, Naomi Taylor, Valérie S. Zimmermann

## Abstract

T cell deficiencies are commonly treated by intravenous hematopoietic stem/progenitor cell (HSPC) transplantation. However, this approach often leads to delayed and incomplete T cell reconstitution, partly due to impaired thymic function. In contrast, intrathymic delivery of HSPCs enables rapid, thymus-autonomous T cell development. Using a ZAP-70-deficient mouse model of severe immunodeficiency, we show that intrathymic transplantation of wild-type HSPCs results in robust thymic engraftment, with mature T cells and FOXP3+ regulatory T cells (Treg) detectable within four weeks. This reconstitution parallels the regeneration of a functional thymic medulla and the emergence of mature medullary thymic epithelial cells (mTECs). Importantly, we identify an early wave of donor-derived RORγT+ ILC3s that correlates with medullary regeneration. While dispensable for steady-state thymopoiesis, ILC3s accelerate medulla formation and support optimal T cell differentiation in this immunodeficient setting. These findings uncover a critical role for ILC3–TEC crosstalk in thymic repair and highlight intrathymic HSPC transplantation as a strategy to enhance immune reconstitution in immunodeficient hosts.

## INTRODUCTION

The thymus is the primary lymphoid organ supporting T cell development, with thymopoiesis initiated upon migration of hematopoietic progenitors from the bone marrow. The thymus is composed of two main zones, the cortex and the medulla, which play essential roles in the different steps of T cell development. In general, cortical thymic epithelial cells (cTECs) promote the commitment of thymic progenitors to the T lineage and ensure the positive selection of thymocytes while medullary thymic epithelial cells (mTECs) and dendritic cells play crucial roles in the negative selection of thymocytes and the generation of suppressive regulatory T cells (Treg) (Kadouri et al., 2020; Ashby and Hogquist, 2024). Indeed, the unique cellular composition and spatial organization of the thymus are essential for the development of a diverse MHC-restricted and self-tolerant T cell repertoire. Notably though, thymocytes reciprocally support the differentiation of cTEC and mTEC through thymic crosstalk (Anderson et al., 2024).

The thymic environment is highly dynamic, evolving during prenatal as well as postnatal periods. During fetal development and early postnatally, the thymic stroma is enriched in cTECs, with lymphoid tissue inducers (LTi) and vγ5^+^ dendritic epidermal T cells (DETC) progenitors—which both express the ligand for the cytokine receptor RANK (RANKL)—playing essential roles in mTEC proliferation and maturation in a RANK-dependent manner (Roberts et al., 2012; Rossi et al., 2007). Interestingly though, in adulthood, immature and mature thymocytes expressing RANKL regulate the homeostasis of the stromal compartment; indeed, the crosstalk between mature SP4 thymocytes and immature mTECs regulates the differentiation state of the latter through RANK-RANKL and CD40-CD40L interactions (van Ewijk et al., 1994; Irla et al., 2012). Nonetheless, with age, the thymus involutes, a process controlled by sex hormones, cytokines, miRNAs, and transcription factors, amongst others (Liang et al., 2022; Li and Zúñiga-Pflücker, 2023; Kousa et al., 2024). Beyond age-related involution, the thymus can undergo physiological or pathological shrinkage in response to stressors such as pregnancy, obesity, infection, or radiotherapy and chemotherapy (Kinsella and Dudakov, 2020). Remarkably, it retains a robust capacity for regeneration, mediated through multiple mechanisms (Anderson et al., 2024; Dudakov and van den Brink, 2025).

Notably, thymus regeneration has recently been shown to be regulated by BMP4 and amphiregulin secreted by endothelial cells and Tregs, respectively (Wertheimer et al., 2018; Lemarquis et al., 2025) as well as by type 2 innate immune responses—with eosinophils (Cosway et al., 2022), tuft mTECs (Nevo et al., 2024), and the IL-25 and IL-33 cytokines all stimulating type 2 innate lymphoid cells (ILC2) (Cosway et al., 2023). Furthermore, type 3 innate immune responses promote thymic regeneration, in large part due to IL-22 production (Dudakov et al., 2012) by type 3 ILC (ILC3) (Dudakov et al., 2017). Lymphotoxin α signaling through the lymphotoxin β receptor (LTβR) (Borelli and Irla, 2021) as well as RANKL administration (Hikosaka et al., 2008; Lopes et al., 2017) also improve thymic regeneration. However, the determinants governing the relative roles of these pathways and their potential interactions remain unclear. Moreover, the mechanisms governing postnatal thymic repair in the setting of genetic defects impairing T cell development remains poorly understood.

Severe combined immunodeficiencies (SCID) represent a heterogeneous group of rare genetic disorders characterized by defects in T cell development, with additional lymphoid lineages affected in some patients (Dvorak et al., 2023). The absence of mature thymocytes disrupts crosstalk with thymic stroma cells, resulting in an abnormal thymic architecture that primarily affects the medulla (van der Burg and van Zelm, 2014; Rucci et al., 2011). If left untreated, these pathologies are generally fatal in childhood. Current treatment options include allogeneic hematopoietic stem cell transplantation (HSCT) or transplantation of autologous gene-corrected HSCs (Dvorak et al., 2023; Fischer and Hacein-Bey-Abina, 2020). Although effective, these therapies are still associated with complications often linked to impaired thymic function, such as prolonged delays in polyclonal T cell reconstitution (>100 days) and the risk of graft-versus-host disease following non-histocompatible transplants (Saglio et al., 2015; Lankester et al., 2022; Eissa et al., 2024). Enhancing thymic function following HSCT could therefore accelerate T cell recovery and improve patient outcomes.

Our group has previously demonstrated the potential of directly targeting hematopoietic stem/ progenitor cells (HSPCs) into the thymus using a ZAP-70-deficient mouse model of SCID. Intrathymic transplantation results in a more diverse T-cell reconstitution, requiring approximately 10-fold fewer HSPCs compared with conventional intravenous HSCT (Adjali et al., 2005b; de Barros et al., 2013). Furthermore, intrathymic HSCT supports long term, self-autonomous thymopoiesis and is associated with restoration of the thymic architecture (Vicente et al., 2010; Peaudecerf et al., 2012; Martins et al., 2012). However, the mechanisms underlying thymic reconstitution, and particularly thymic architecture repair, remain unclear.

In this study, we characterize the early steps of T cell development and thymic architecture repair following intrathymic administration of WT HSPCs into ZAP-70-deficient mice, and identify the mechanisms driving rapid thymic restoration. Intrathymically administered WT HSPCs efficiently engraft and sustain long-term T cell development. Thymic medulla formation, observed 3 weeks post-transplantation, was accompanied by substantial differentiation of donor-derived RORγT⁺ ILC3. While ILCs are dispensable for physiological thymopoiesis, as demonstrated in *Nfil3*^-/-^ mice lacking ILC progenitors (Xu et al., 2015), we reveal their critical role in thymic repair and long-term T cell development under conditions of immunodeficiency.

## RESULTS

### Rapid engraftment and expansion of WT BM progenitors following intrathymic transplantation into immunodeficient *Zap-70*^-/-^ mice

To evaluate the impact of IT transplantation on the kinetics of expansion of donor HSPC in the context of an immunodeficient thymus, we took advantage of the ZAP-70–deficient mouse model exhibiting an arrest in thymocyte differentiation at the DP stage (Au-Yeung et al., 2018; Negishi et al., 1995). Following IT administration of BM HSPCs (2×10^5^ lineage-negative (lin^−^) CD45.1^+^ cells) into non-conditioned 2.5-week-old *Zap-70*^−/−^ mice (CD45.2^+^) (**Figure 1A)**, we observed a rapid increase in the percentages of donor cells between 2- and 3-weeks post-transplant (0.2±0.2% vs 12.7±8%, respectively; p<0.001) and these percentages of donor cells were maintained at a plateau from 4 to >15w post-transplant (19.4±4% vs 25.1±13%, respectively; **Figures 1B-C**). While this increase represents an approximately 100-fold expansion of donor progenitors, it is notable that the total number of thymocytes remains stable following IT HSPC transplantation (**Figure 1C**). Importantly, no significant thymic engraftment or expansion of donor cells was observed following IV transplantation of HSPC into non-conditioned *Zap-70*^-/-^ mice (**Figures S1A and S1B**).

**Figure 1:**
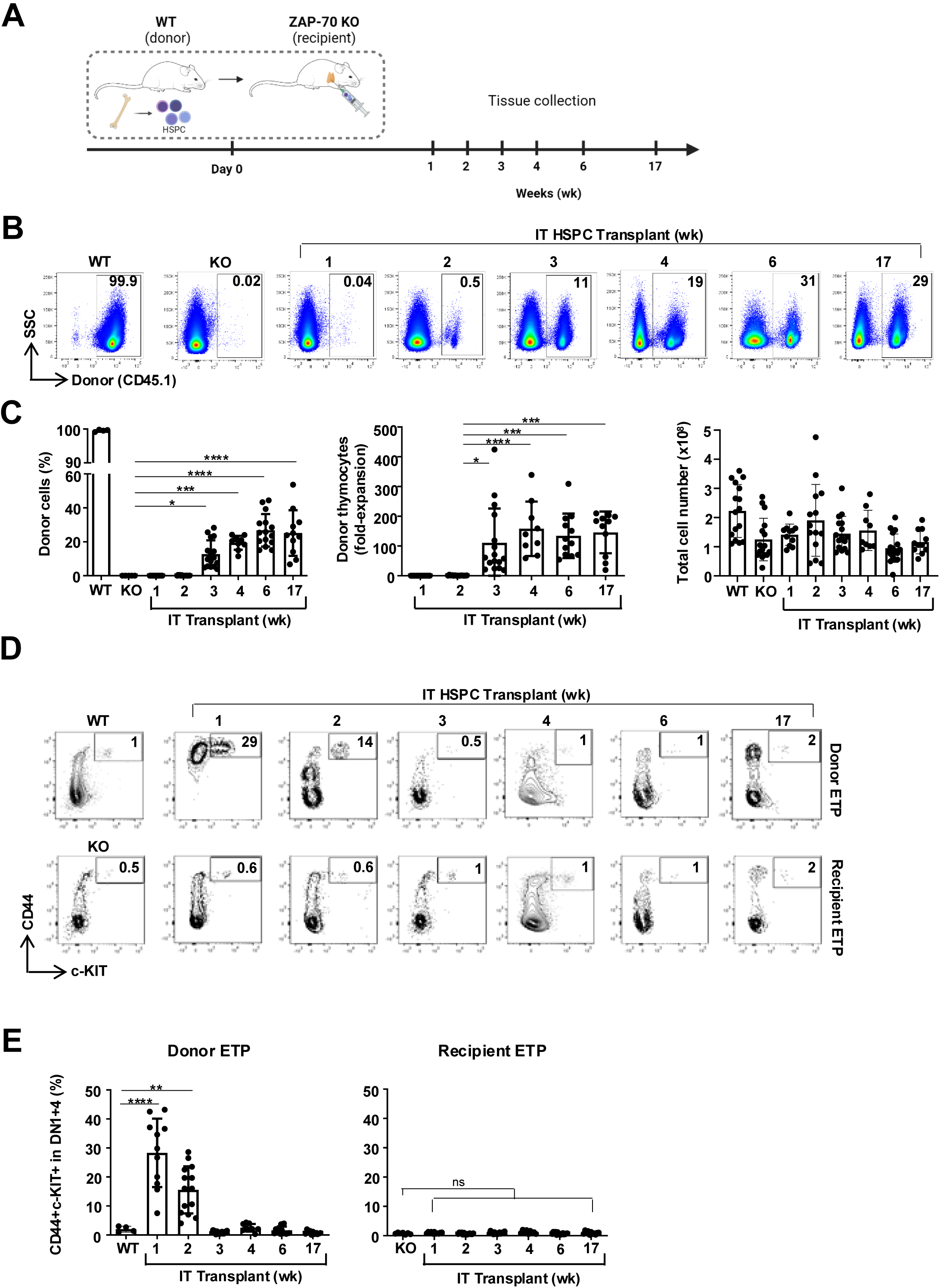
Rapid engraftment and expansion of donor HSCs following intrathymic administration into *Zap-70*^−/−^ mice. **(A)** Non-conditioned *Zap-70*^−/−^ (KO) mice (CD45.2^+^) were transplanted by intrathymic administration of WT histocompatible BM progenitors (CD45.1^+^, 2×10^5^). Mice were sacrificed at weeks 1, 2, 3, 4, 6 and 17 post-transplant. Thymus and lymph nodes were harvested and analyzed by flow cytometry. **(B)** The presence of CD45.1^+^ donor cells was monitored by flow cytometry. Representative dot plots of thymi at 1, 2, 3, 4, 6 and 17 weeks after transplantation are presented and the percentages of CD45.1^+^ cells are indicated. Controls showing CD45.1^+^ thymocytes in WT and KO recipients are also presented. **(C)** Quantification of the percentages of CD45.1^+^ donor cells in the thymi of *Zap-70*^-/-^mice (left), fold-expansion relative to the number of injected progenitors (middle), and total thymocyte numbers (right) are presented at 1-17 weeks post transplantation. Each point represents data from an individual thymus. **(D)** Representative dot plots showing the percentages of donor (upper panel) and recipient (lower panel) early thymic progenitors (ETP) at the indicated time points following IT transplantation, evaluated as a function of CD44/c-KIT expression in CD25^-^ double negative (DN) thymocytes. **(E)** Quantification of the percentages of donor (left) and recipient (right) ETPs at the indicated time points (n=8-16 mice per time point) are presented. Means ± SD are shown and data were obtained in at least 2 independent experiments. Statistical significance was determined using a 1-way ANOVA with a Tukey multiple comparison test. *p<0.05; **p<0.01; ***p<0.001; ****p<0.0001

Donor HSPC engraftment was associated with an expansion of early thymic progenitors (ETP; CD44^+^C-KIT^+^ double negative (DN) thymocytes) within 1 week following IT transplantation (28±12%). This level of ETPs was significantly higher than that detected in WT mice (2±1%; p<0.0001), highlighting a selective advantage of donor ETP at a stage of differentiation wherein the ZAP-70 kinase is not expressed (Palacios and Weiss, 2007). Notably though, ETP levels decreased to those detected in the WT thymus by 3 weeks post IT transplantation and remained stable over the next 4 months (0.8±0.6%; **Figures 1D-E**). Thus, our data show that WT HSPCs exhibit a rapid engraftment and selective advantage following their IT administration in *Zap-70*^−/−^ mice with an early expansion of donor ETP and a long-term maintenance of donor thymocytes.

### IT-transplanted BM progenitors rapidly generate mature thymocytes and support long-term T cell differentiation

To determine the differentiation state of engrafted donor progenitors as a function of time post IT transplantation, CD4/CD8 thymocyte profiles were evaluated at different time points. At 1-2 weeks post-transplantation, high levels of immature double negative (DN) thymocytes were detected (43±12% and 42±6, respectively), consistent with early stages of thymopoiesis, and by week 3, they had returned to baseline levels (3.9±2%). Progression to the double positive (DP) thymocyte stage increased between weeks 2 and 3 post transplantation, from 50±7% to 88±2% (p<0.0001, **Figures 2A-B**). Importantly, thymopoiesis was then maintained as the percentages of DP thymocytes remained constant from week 3, with levels similar to those detected in WT mice throughout the 17-week experimental period. Furthermore, percentages of CD4^+^CD8^-^ and CD8^+^CD4^-^ thymocytes increased, reaching a maximum at 4 weeks post transplantation (14±3% and 6±2%, respectively, **Figures 2A-B**). To evaluate the differentiation state of these thymocytes, CD3 expression was monitored; at 2 weeks post IT transplantation, the vast majority of CD8^+^CD4^-^ cells were CD3^-^ (92±9%), reflecting immature single positive CD8 cells (ISP8; **Figure 2C**), but CD3 expression increased dramatically between weeks 3 and 4 of differentiation (47±15% to 87±8%), reaching levels detected in WT mice (**Figures 2C-D**). Of note, CD3^+^ mature SP4 thymocytes were detected at an earlier time point than SP8 thymocytes, by 3 weeks post IT transplantation (**Figure 2C-D**). Interestingly, although SP4 thymocytes do not normally pass through a CD3^-^ intermediate stage during physiological differentiation, we were surprised to detect a substantial population of CD3^-^CD4^+^ thymocytes at this early time point, comprising 76±25% and 24±16% of cells at 2 and 3 weeks, respectively. Nonetheless, the percentages of CD3^+^CD4^+^ cells rapidly progressed, accounting for 76±16% and 85±12% of all SP4 thymocytes by weeks 3 and 4 of differentiation (**Figure 2C-D**). This differentiation led to the export of mature donor-derived T cells into the periphery, accounting for 10±2.5% of all lymph node cells at week 4 and rising to 34±4% by 17 weeks post-transplantation (**Figures 2E-F**). These peripheral donor cells represented >90% CD4 and CD8 T cells, with a CD4/CD8 ratio similar to that detected in WT mice (**Figures S2A-B**), highlighting the robust thymopoiesis in these mice. This differentiation led to the export of mature donor-derived T cells into the periphery, accounting for 10±2.5% of all lymph node cells at week 4 and rising to 34±4% by 17 weeks post-transplantation (**Figures 2E-F**). The vast majority of donor-derived peripheral cells were T cells, with a CD4/CD8 ratio similar to WT mice (**Figures S2A-B**), underscoring the robustness of thymopoiesis in this setting.

**Figure 2:**
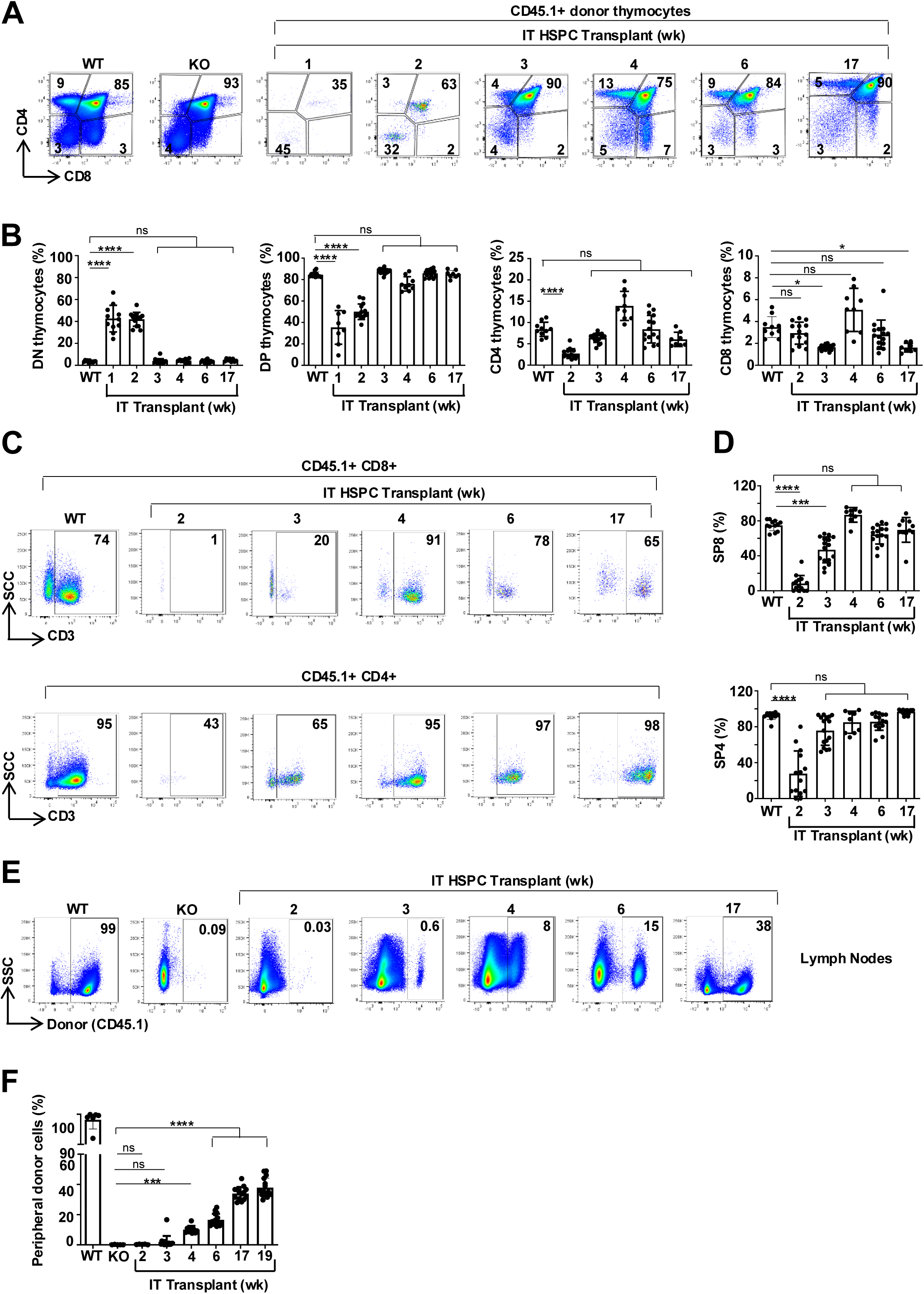
Rapid generation of mature thymocytes following intrathymic HSC transplantation. **(A)** Representative CD4/CD8 profiles of thymi from WT, *Zap-70*^−/−^ (KO) and IT-transplanted KO mice are shown at the indicated time points and the percentages of DN, DP, CD4, and CD8 thymocytes are noted. **(B)** Quantification of the percentages of DN, DP, CD4 and CD8 donor cells detected in the thymi of IT-reconstituted *Zap-70*^−/−^ mice are presented at the indicated time points and each point presents data from an individual mouse thymus (n= 8-16 per group from at least 2 independent experiments). **(C)** Representative plots showing expression of CD3 within CD8^+^ (upper plots) and CD4^+^ (lower plots) thymocytes are presented at each time point. **(D)** Quantification of the percentages of CD3^+^ cells within the CD8 (upper panel) and CD4 (lower panel) thymocyte subsets are presented for individual mice between 2-17 weeks post transplantation (n= 8-16 per group from 2 independent experiments). **(E)** Representative plots showing the percentages of CD45.1^+^ cells in lymph nodes (LN) of WT and IT-transplanted mice at 2-17 weeks post transplantation. **(F)** Quantification of CD45.1^+^ cells in LNs of mice at 2-17 weeks post-transplantation and means ± SD are presented. Statistical significance was determined using a 1-way ANOVA with a Tukey multiple comparison test. ns, not significant; *p<0.05; **p<0.01; ***p<0.001; ****p<0.0001

### IT BM progenitor transplantation results in a rapid generation of FOXP3^+^ Treg cells

As the differentiation of FOXP3^+^ Treg in the thymus has been associated with improved patient outcome following allogeneic HSCT (Edinger and Hoffmann, 2011), we next evaluated the kinetics of differentiation of FOXP3^+^ SP4 Treg following IT progenitor transplantation in this immunodeficient setting. To study thymic Treg differentiation, we monitored the presence of CD25^+^FOXP3^-^ and CD25^-^FOXP3^+^ SP4 thymocytes, 2 progenitor populations that have been reported to give rise to mature CD25^+^FOXP3^+^ Treg (Owen et al., 2019a; Tai and Singer, 2014) (**Figure 3A)**. Interestingly, CD25^+^FOXP3^-^ progenitors developed prior to CD25^-^FOXP3^+^ progenitors (**Figures 3B-C**), consistent with recent work highlighting the presence of agonist-signaled CD25^+^ precursors (Tai et al., 2023; Tai and Singer, 2024). The presence of these CD25^+^FOXP3^-^ progenitors reflected a high level of Treg differentiation at early time points post transplantation as these cells were present at levels significantly higher than detected in the WT thymus (2.0±0.5% vs. 0.9±0.3%; p<0.0001). By 4 weeks post differentiation, there was an increase in CD25^-^FOXP3^+^ progenitors, to levels similar to that detected in WT mice (1.2±0.3% and 1.3±0.6%, respectively, **Figures 3B-C**). The presence of this subset as well as the more mature CD25^+^FOXP3^+^ Treg subset at 4 weeks post IT transplantation correlated with the export of mature Treg to the periphery. Indeed, CD25^+^FOXP3^+^ Treg accounted for a similar percentage of peripheral CD4 LN T cells at 4 weeks in IT-transplanted and WT mice (14±8% and 15±2%, respectively; **Figures 3D-E**). Together, these data suggest that Treg differentiation from IT-transplanted progenitors through an initial CD25^+^FOXP3^-^ stage, followed approximately 1 week later by the emergence of CD25^-^FOXP3^+^ and CD25^+^FOXP3⁺ cells, coinciding with the appearance of CD3^+^ SP4 thymocytes (**Figures 2D** - **3C**).

**Figure 3:**
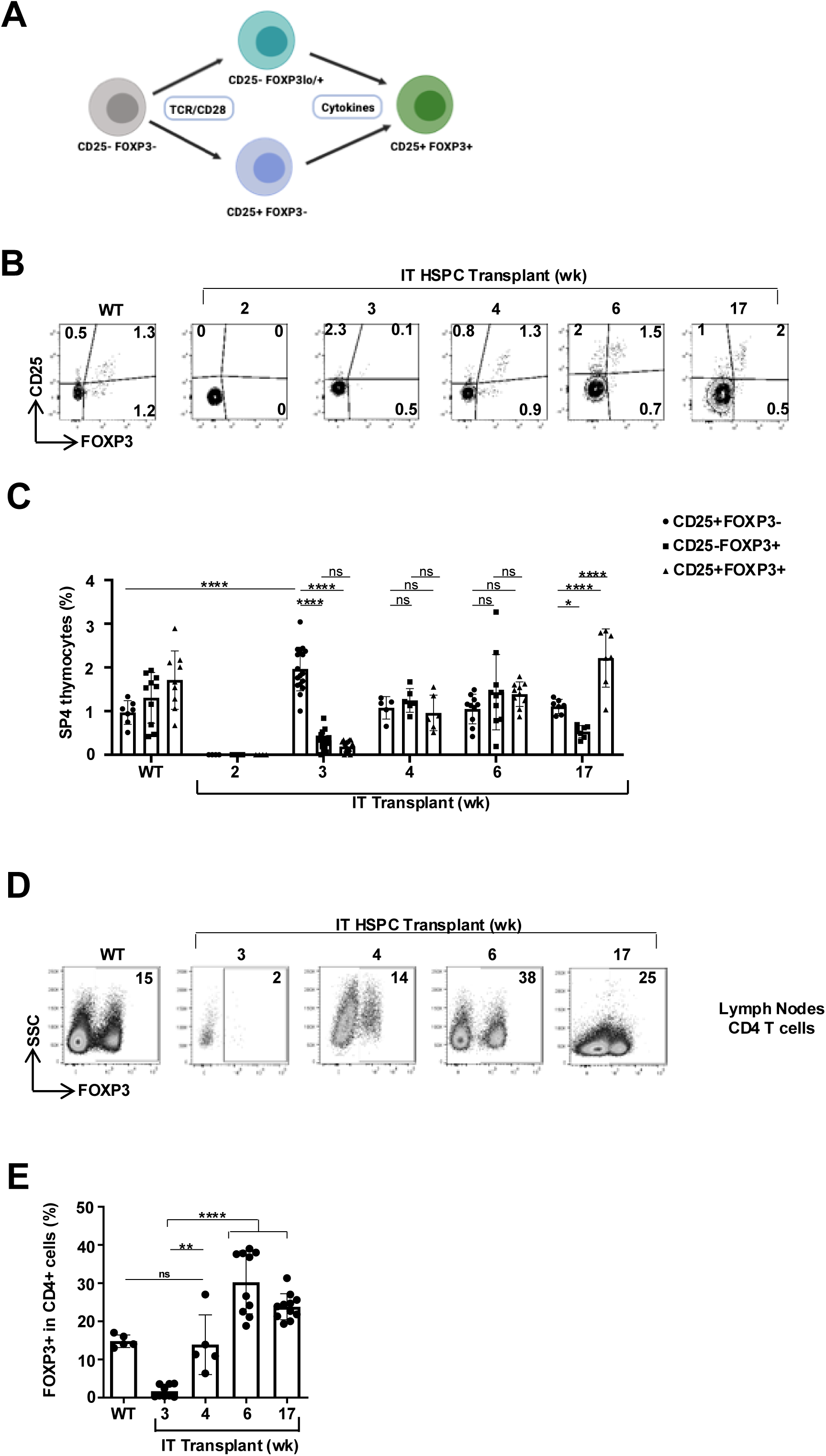
Kinetics of thymic Treg differentiation following IT HSPC transplantation. **(A)** Schematic representation of thymic regulatory T cell differentiation models **(B)** Differentiation of thymic Treg was evaluated as a function of CD25/FOXP3 profiles within SP4 thymocytes and representative plots in WT mice and following IT transplantation of *Zap-70*^−/−^ mice are presented. **(C)** Quantification of CD25⁺FOXP3⁻, CD25⁻FOXP3⁺, and CD25⁺FOXP3⁺ thymocytes (n = 5–17 per group) was performed across two independent experiments from weeks 2 to 17. Data are shown as means ± SD. **(D)** Expression of FOXP3 in LN CD4^+^ cells was evaluated by flow cytometry and representative plots are presented. **(E)** The percentages of Foxp3^+^ cells within LN CD4 lymphocytes were evaluated at the indicated time points and means ± SD are presented (n=5-11 per group). Statistical significance was determined using a 1-way ANOVA with a Tukey multiple comparison test. *p<0.05; **p<0.01; ***p<0.001

### IT transplantation of WT BM progenitors leads to the rapid generation of medullary thymic epithelial cells

Treg differentiation is dependent on the presence of a functional thymic medulla (Perry et al., 2014) and this structure is defective in the ZAP-70-deficient thymus (Negishi et al., 1995; Pouzolles et al., 2020). Medulla formation is associated with the differentiation of AIRE-expressing mTECs, an epithelial subset not present in immunodeficient thymi (Rucci et al., 2011) and dependent on the crosstalk between TEC and mature thymocytes (Desanti et al., 2012). To evaluate changes in the thymic architecture following IT HSPC transplantation, K14^+^ mTEC zones and AIRE^+^ mTEC were monitored on thymus slices as a function of time post transplantation. By 3 weeks post-transplantation, the number of large K14^+^ mTEC zones (>10 000 μm^2^) increased significantly as compared to ZAP-70 deficient mice (3±2.6 to 13±8, respectively; p<0.001, **Figures 4A-B**) and remained stable through week 17 (14±4). Similarly, AIRE^+^ cell numbers rose markedly by week 4 (from 20±13 to 99±35) and remained elevated for the duration of the 17-week experimental period (93±29/mm^2^, p<0.0001; **Figures 4A-B**). These data suggest that a rapid expansion of mTEC occurs at approximately 3 weeks post IT transplantation followed by their maturation to AIRE-expressing mTEC by 4 weeks. Importantly though, medulla formation was not significantly modulated by IV HSPC transplantation, at time points ranging from 4-17 weeks post transplantation (**Figures S3A-B**). Thus, intrathymic HSPC transplantation, unlike intravenous delivery, effectively promoted the generation of a physiological thymic architecture.

**Figure 4.**
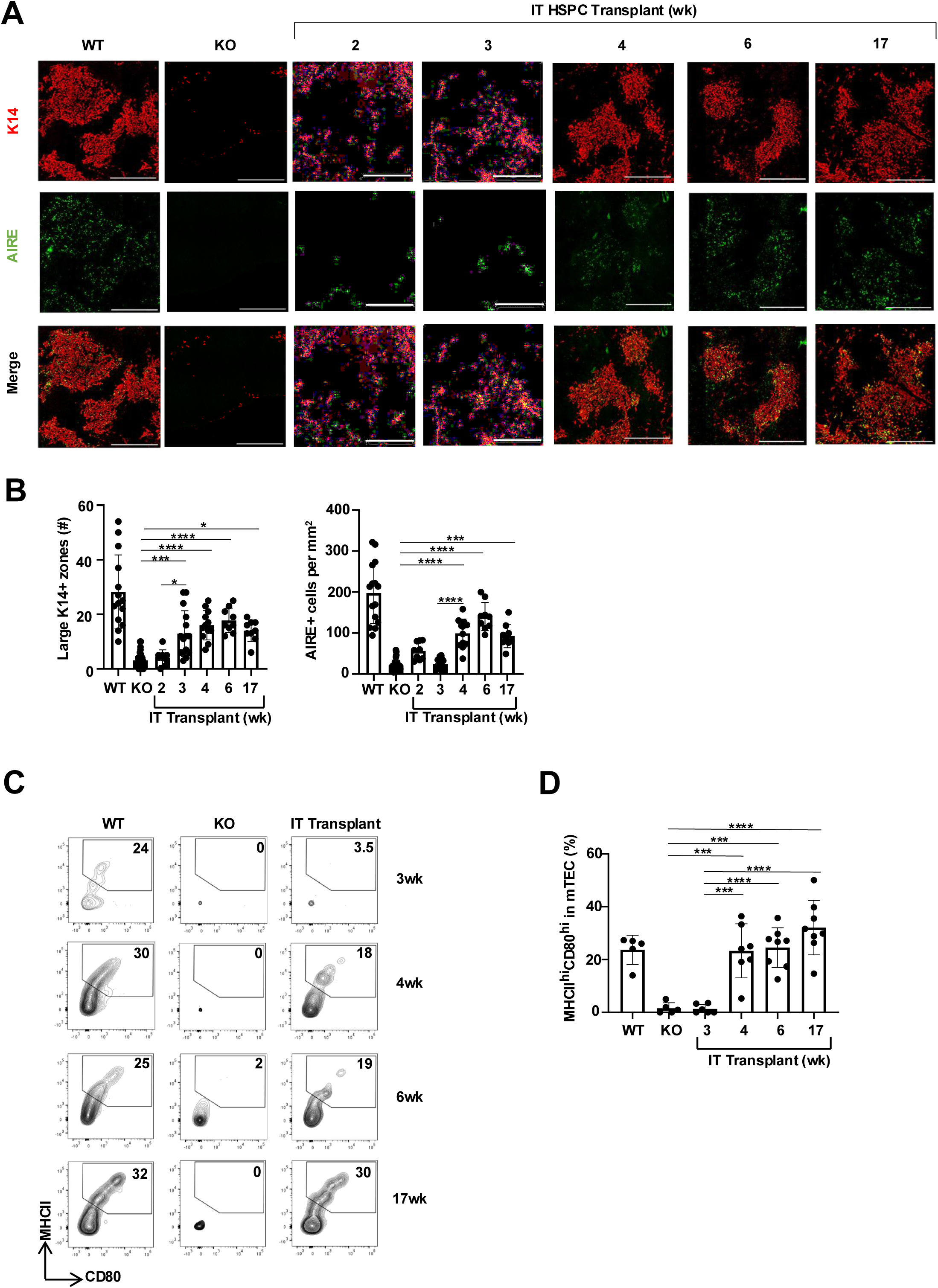
IT HSPC transplantations result in a rapid restoration of the thymic medulla. **(A)** Thymi from WT, KO, and transplanted mice were evaluated for the presence of a thymic medulla by immunohistochemical analysis of keratin 14 (K14) and AIRE expression. Representative microscopy images of thymic tissue sections at 2, 3, 4, 6 and 17 weeks after IT injection of HSPCs are shown. **(B)** Quantification of the numbers of large K14^+^ zones (>10 000/ mm^2^; left panel) and the number of AIRE^+^ cells per mm^2^ (right panel) are presented for the indicated conditions. Quantifications were performed in medulla thymus from at least 3 individual mice. **(C)** The presence of CD80^hi^MHCII^hi^ mTEC was evaluated by flow cytometry following digestion of thymi from WT, *Zap-70*^−/−^, and IT-transplanted *Zap-70*^−/−^ mice, at the indicated time points. mTECs were evaluated on gated EPCAM^+^UEA-1^+^ cells and representative plots are shown. **(D)** Quantification of CD80^hi^MHCII^hi^ mTEC in WT and *Zap-70*^−/−^ mice at the indicated time points after IT HSPC transplantation (n=5-8 mice per group). Means ± SD are presented. Statistical significance was determined using a 1-way ANOVA with a Tukey multiple comparison test. ns, not significant; *p<0.05; **p<0.01; ***p<0.001; ****p<0.0001

To specifically monitor the mTEC maturation, we assessed expression of CD80 and MHCII, as previously described (Bornstein et al., 2018; Miragaia et al., 2018). Consistent with the data above, mature CD80^hi^MHCII^hi^ UEA-1^+^ mTECs appeared by 4 weeks post-transplantation, increasing from 1.5±2% to 23±10% (p=0.0004, **Figures 4C-D**), a level comparable to that of WT thymus (24±6%). Notably, mTEC maturation was not observed following IV HSPC transplantation, consistent with the absence of mTEC zones in this setting (**Figures S3C-D**). Together, these findings demonstrate that the emergence of mature mTEC is tightly linked to the first wave of T cell differentiation following IT HSPC transplantation.

### The generation of donor-derived thymic innate lymphoid cells (ILC) is critical for mTEC regeneration and thymocyte differentiation

Intrathymic transplantation of HSPC led to the appearance of a CD3^-^CD4^+^ thymocyte subset (**Figures 2C-D**), which is generally not observed during the differentiation of mature SP4 thymocytes. Quantification revealed that CD3^-^CD4^+^ donor cells accounted for 76±25% of all CD4^+^ thymocytes at 2 weeks post-transplantation, an 8-fold increase compared with WT thymus (p<0.0001, **Figure 5A**). To further characterize these cells, we performed single cell RNAseq on FACS-sorted CD3^-^CD8^-^ thymocytes at 3 weeks post IT HSPC transplantation, distinguishing recipient and donor populations. Both immature double-negative thymocytes and ILCs were detected (**Figures 5B** and **S4A-B**). Significant differences were observed in the relative frequencies of recipient versus donor thymocytes (**Figures S4C-D**), with donor cells notably enriched for type 3 ILCs (ILC3) and lymphoid tissue inducer (LTi) cells, a subset of ILC3 (**Figure 5C**).

**Figure 5.**
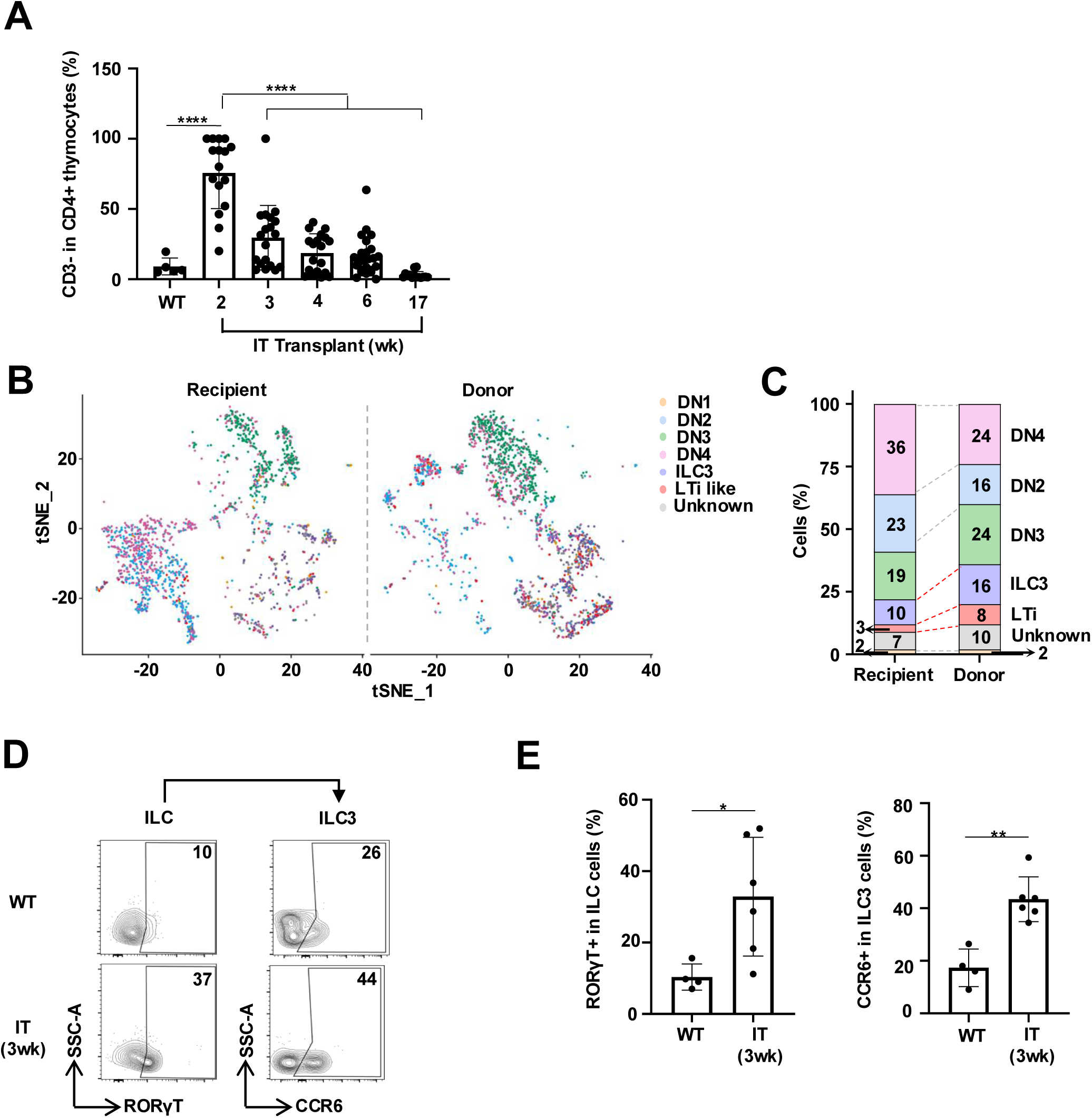
IT HSPC transplantation promotes the development of an ILC3-like population. **(A)** Percentages of CD3^-^ cells in CD8^-^CD4^+^ thymocytes at different time points post IT-HSPC transplantation are depicted. **(B)** CD3^-^CD8^-^ thymocytes were FACS-sorted from *Zap-70*^−/−^ mice at 3 weeks following IT HSPC transplantation and subjected to single-cell RNAseq. Spectral tSNE plots of recipient (left) and donor (right) cells colored by density clustering and annotated by cell-type identity are presented. **(C)** The percentages of double negative (DN1, DN2, DN3, and DN4), type 3 innate lymphoid cells (ILC3), and lymphoid tissue inducer (LTi), an ILC3 subset, within recipient (left) and donor (right) thymocytes are presented. **(D)** The presence of RORγT^+^ ILC3 were evaluated on gated CD45.1^+^CD8^-^TCRβ^-^TCRγδ^-^CD4^+^CD127^+^ cells and representative dot plots in WT and IT-transplanted *Zap-70*^−/−^ mice are shown (left panels). Expression of CCR6 within the RORγT^+^ ILC3 subset are presented (right panels). **(E)** Quantification of the percentages of RORγT^+^ ILC3 and CCR6^+^ ILC3 (LTi) in the thymi of WT and IT-transplanted *Zap-70*^−/−^ mice (n=4 for WT and n=6 for IT-transplanted). Means ± SD are presented. Statistical significance was determined using a 1-way ANOVA with a Tukey multiple comparison test. ns, not significant; *p<0.05; **p<0.01

To confirm ILC3 identity, we assessed RORγT expression within the ILC subset, a marker specific for ILC3 (Scoville et al., 2019; Vivier et al., 2018). Notably, at 3 weeks post-transplantation, donor thymocytes contained a significantly higher percentage of RORγT^+^ ILC3s than WT thymus (33±17% vs 10±4%, respectively; p<0.05, **Figure 5D-E**). Furthermore, the LTi subset expressing CCR6 was enriched in donor RORγT^+^ ILC3s compared with WT (43±8% vs 17±7%, respectively; p<0.05, **Figures 5D-E**). These results indicate that robust thymopoiesis following IT HSPC transplantation is associated with an early and substantial increase in donor RORγT^+^ ILC3s.

### The generation of donor-derived thymic innate lymphoid cells (ILC) is critical for mTEC and thymocyte differentiation

The ILC3 subset is not required for physiological postnatal thymus differentiation, but it positively regulates mTEC generation and thymopoiesis under pathological conditions via IL-22 production and RANKL expression (Dudakov et al., 2012; Lopes et al., 2017). We therefore asked whether donor-derived ILCs, and particularly ILC3s, are required for thymopoiesis following IT HSPC transplantation in immunodeficient mice. To test this, ZAP-70-deficient mice were IT-transplanted with HSPCs lacking the NFIL3 transcription factor (**Figure 6A**). NFIL3 has been shown to be essential for ILC development (Seillet et al., 2014) and *Nfil3*^-/-^ mice lack ILC progenitors (Xu et al., 2015; Léger et al., 2025). As expected, the absence of *Nfil3* did not alter the thymic architecture and thymopoiesis in physiological conditions (Kashiwada et al., 2010), and we observed no significant differences in large K14+ mTEC zones or AIRE^+^ cells between *Nfil3*^+/+^ and *Nfil3*^-/-^ mice (**Figures 6B-C**).

**Figure 6.**
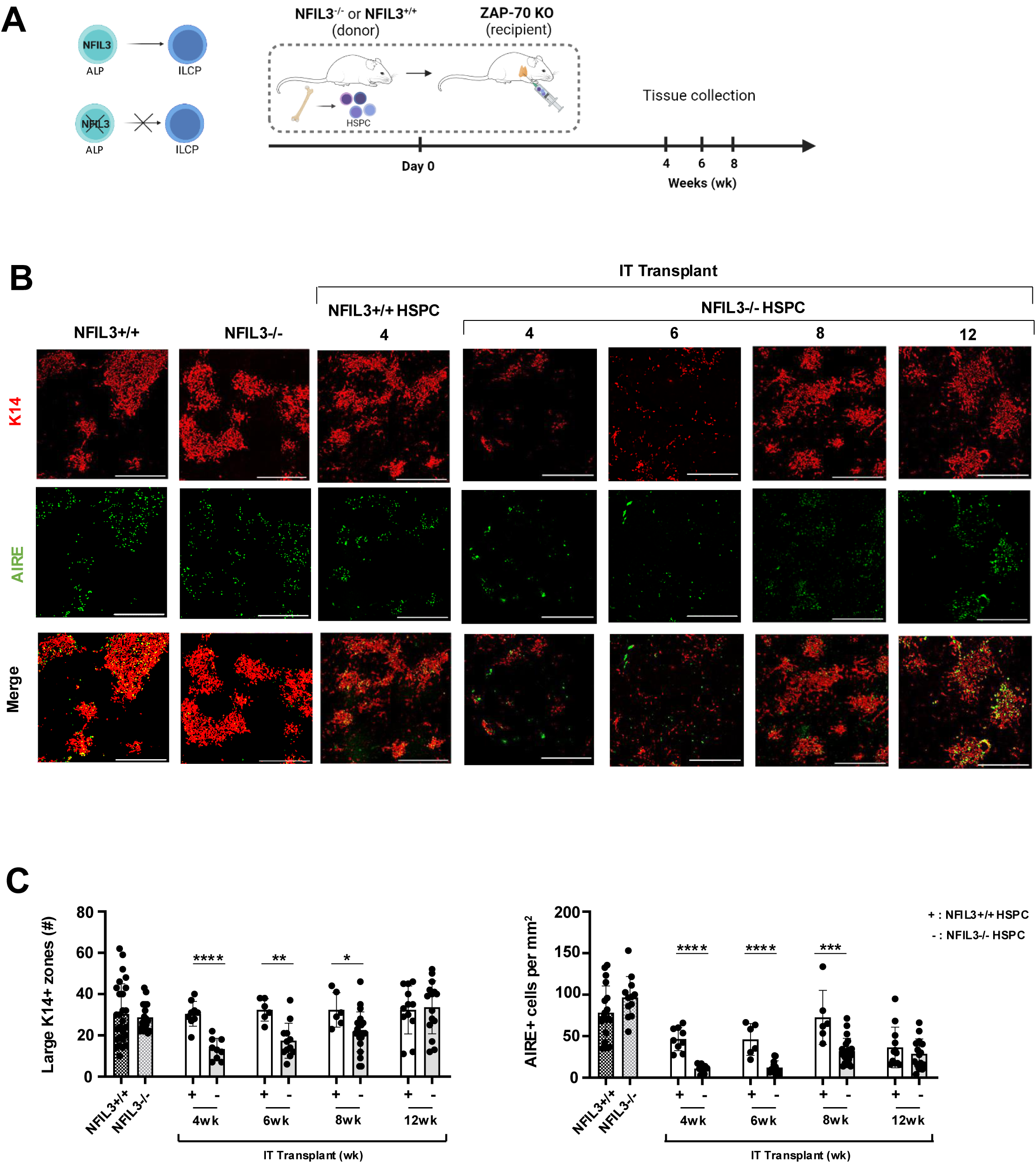
The generation of thymic ILCs is critical for rapid medulla generation. **(A)** Schematic of the IT transplantation of *Zap-70*^−/−^ mice with *Nfil3*^+/+^ or *Nfil3*^−/−^ BM progenitors (2×10^5^). Mice were sacrificed at 4, 6, 8 and 12 weeks post-transplant. Thymus were harvested and analyzed by flow cytometry or by immunochemistry. **(B)** The presence of a thymic medulla was evaluated by immunohistochemical analysis of keratin 14 (K14) and AIRE expression and representative microscopy images of thymic tissue sections are shown. **(C)** Quantification of the numbers of large K14^+^ zones (>10 000/ mm^2^; left panel) and the number of AIRE^+^ cells per mm^2^ (right panel) are presented for the indicated conditions. Quantifications were performed in medulla thymi from at least 3 individual mice from 2 different experiments. Statistical significance was determined using an unpaired 2-tailed t-test. ns, not significant; *p<0.05; **p<0.01; ***p<0.001; ****p<0.0001

Notably though, in the context of IT transplantation into *Zap-70*^-/-^ mice, recipients of *Nfil3*^-/-^ progenitors exhibited delayed medulla generation compared with recipients of *Nfil3*^+/+^ progenitors. The former presented with significantly fewer large K14^+^ mTEC zones and AIRE^+^ mTECs at 4-8 weeks post transplantation (30±6 vs 13±5 and 47±5 vs 11±2, at 4w respectively; p<0.05; **Figures 6B-C and Figure S5**). By 12 weeks, medulla generation reached similar levels in both groups. These results demonstrate the crucial role of donor derived-ILC in the rapid generation of the thymic medulla in the immunodeficient thymus.

Although the initial delay in mTEC generation did not impact the early engraftment and expansion of donor thymocytes in *Zap70*^-/-^ mice, by 8 weeks there were significantly fewer donor thymocytes in mice transplanted with *Nfil3*^-/-^ progenitors (17±6% vs 5±7%, respectively; p<0.001; **Figure 7A**). This reduction was associated with a loss of ongoing thymopoiesis, defined as >50% DP thymocytes (**Figure 7B**), and a decrease in donor ETP (**Figure 7C**).

**Figure 7.**
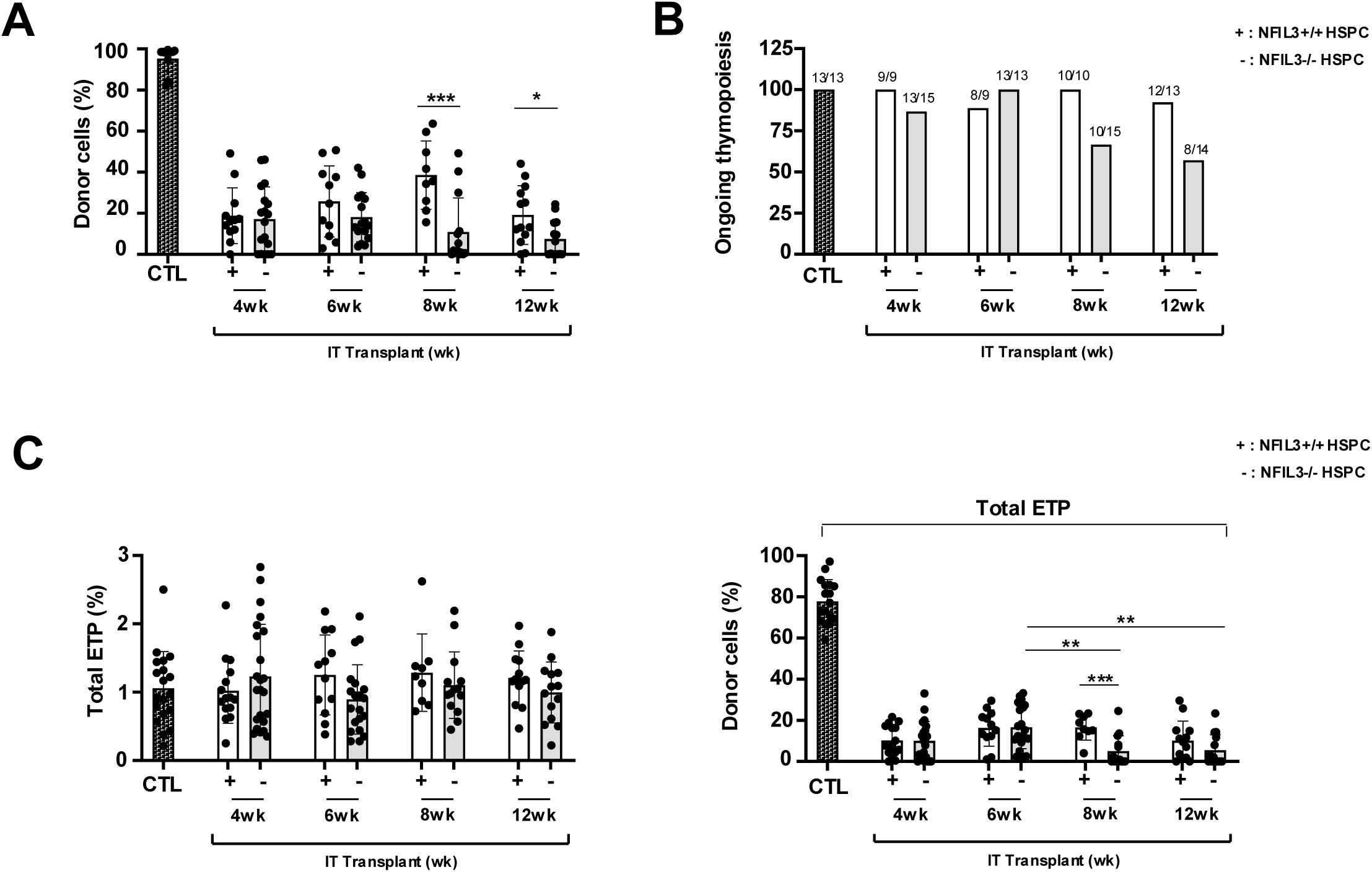
Donor-derived thymic ILCs are critical for long term thymopoieisis maintenance following IT HSPCT. **(A)** IT transplantation of *Zap-70*^−/−^ was performed as described in Figure 6A. Quantification of YFP^+^ donor thymocytes at 4, 6, 8 and 12 weeks following IT administration of *Nfil3*^+/+^ or *Nfil3*^−/−^ HSPCs into *Zap-70*^−/−^ mice. Means ± SD are presented. **(B)** Representation of ongoing thymopoiesis evaluated as DP >50%– at the different time points following IT administration of *Nfil3*^+/+^ or *Nfil3*^−/−^ HSPCs into *Zap-70*^−/−^ mice. The number of mice analyzed is specified on each bars **(C)** Quantification of the percentages of total ETP (left) and donor cells within the ETP population (right) at 4, 6, 8 and 12 weeks following IT administration of *Nfil3*^+/+^ or *Nfil3*^−/−^ HSPCs in *Zap-70*^−/−^ mice. Statistical significance was determined using a 1-way ANOVA with a Tukey multiple comparison test. ns, not significant; *p<0.05; ***p<0.001; ****p<0.0001

Together these data highlight the importance of intrathymic HSPC transplantation in fostering a reconstitution of the thymic architecture and supporting optimal donor-derived thymopoiesis. They also reveal the critical role of ILC3-TEC crosstalk in promoting immune reconstitution in the setting of genetic immunodeficiencies.

## DISCUSSION

The thymus plays a central role in T cell development and the induction of immune tolerance, sustaining adaptive immunity across the lifespan. While it undergoes natural age-related involution, the thymus is also highly sensitive to stress, including infections, conditioning regimens, or intrinsic genetic deficiencies. Remarkably, it retains a unique regenerative capacity that can be harnessed therapeutically following HSCT or thymic injury (Anderson et al., 2024; van der Burg and van Zelm, 2014; Rucci et al., 2011). However, the mechanisms driving repair in intrinsic thymic deficiencies remain poorly understood.

Here, we demonstrate that intrathymic (IT) transplantation of wild-type HSPCs into ZAP-70-deficient mice enables rapid and sustained thymic reconstitution. By bypassing bone marrow homing, IT delivery allows immediate local seeding of donor cells, accelerating thymopoiesis and enhancing engraftment efficiency (Zlotoff et al., 2010; Schwarz et al., 2007; Foss et al., 2001; Vicente et al., 2010; de Barros et al., 2013). Donor cells progress through canonical thymocyte stages—ETP/DN (1 week), DP (2 weeks), and SP (3–4 weeks)—with CD4 SP thymocytes appearing before CD8 SP cells, establishing continuous thymopoiesis. The kinetics mirror physiological differentiation, indicating a functional niche capable of sustaining progenitor seeding and differentiation (Porritt et al., 2003; Singer et al., 2008; Karimi et al., 2021), and occurs without leukemic transformation (Peaudecerf et al., 2012; Martins et al., 2012).

Thymopoiesis is regulated by reciprocal signaling between thymocytes and thymic epithelial cells (TEC). Following IT transplantation, donor progenitors initiate a synchronized wave of differentiation that coincides with restoration of thymic architecture, particularly the medullary compartment. Within four weeks, immature mTEC^lo^ differentiate into AIRE⁺ mTEC^hi^, reestablishing central tolerance. This regeneration is not observed after intravenous HSPC delivery, reflecting the importance of spatial organization for orderly thymocyte progression. Critically, donor-derived LTi-like RORγt⁺CCR6⁺ ILC3s emerge coincident with medullary recovery. While dispensable for steady-state postnatal thymopoiesis, these cells promote regeneration under immunodeficient or injured conditions (Dudakov et al., 2012; Lopes et al., 2017; Gäbler et al., 2007). Consistent with this, IT transplantation of *Nfil3*-deficient progenitors, which notably lack ILC3, delayed mTEC maturation and ultimately reduced donor thymocyte output, highlighting the essential role of ILC3–TEC crosstalk in establishing a functional niche. Other innate pathways, including ILC2, eosinophils, or tuft cells, were not significantly involved, underscoring ILC3 specificity in this immunodeficient context, though complementary contributions cannot be fully excluded.

Thymic Treg differentiation depends on medullary integrity (Cowan et al., 2015; Irla et al., 2008). In our model, CD25⁺FOXP3⁻ and CD25⁻FOXP3⁺ Treg progenitors emerged sequentially following SP4 thymocyte differentiation and coincided with medullary restoration, highlighting the interplay between thymic architecture and central tolerance. Timely Treg generation is critical for controlling autoimmunity and graft-versus-host disease after HSCT (Edinger and Hoffmann, 2011; Edinger et al., 2003).

In conclusion, intrathymic HSPC transplantation enables rapid and durable thymic reconstitution, restores medullary architecture, and supports functional thymopoiesis through a critical ILC3–TEC axis. These findings emphasize the mechanistic importance of stromal–hematopoietic crosstalk in immune regeneration and provide a foundation for strategies to improve thymic function and immune reconstitution following HSCT or thymic injury.

## MATERIALS AND METHODS

### Mice, cell selections and BM transplantation protocols

*Zap-70*^−/−^ mice (CD45.2^+^) were provided by A. Singer (National Institutes of Health, Bethesda, MD), and the strain was bred and maintained under pathogen-free conditions. Donor BM cells, harvested from the femurs and tibias of either WT CD45.1^+^ C57Bl/6J or *Nfil3^−/−^* mice, generously provided by C. Harly (Gascoyne et al., 2009), were first incubated with a cocktail of rat α mouse antibodies (Abs) directed against lineage markers (CD4, CD8, CD3, TCRγδ [BioXcell, West Lebanon, NH], Ter119, B220, CD11b, and Gr-1 [Thermofisher]) and then with α-rat IgG magnetic beads (Dynal) to deplete differentiated hematopoietic cells. For HSPCT transplantations, 2 × 10^5^ lineage-negative HSPCs were administered either IV or IT into *Zap-70*^−/−^ mice. Intrathymic injections were performed as previously described (Adjali et al., 2005a). For ILC enrichment, CD8^+^ cells were depleted from thymocytes of reconstituted mice, euthanized 3 weeks after transplantation, as well as from age-matched WT control mice. In brief, double-negative (DN) thymocytes and CD4^+^ thymocytes were recovered by depleting CD8^+^ thymocytes using a specific antibody (CD8 rat α mouse hybridoma Abs [BioXcell, West Lebanon, NH]) followed by incubation with α-rat IgG magnetic beads (Dynal). All experiments were approved by the local animal facility institutional review board in accordance with national guidelines.

### Immunophenotyping and flow cytometry analyses

Cells, isolated from thymi, BM or LN, were first stained with a Live/Dead viability dye (ThermoFisher) to exclude non-viable cells, and subsequently stained with the appropriate conjugated αCD45.1 (A20), αc-KIT (2B8), αCD4 (RM4A), αCD8α (53-5.8), αCD25 (PC61), αCD44 (IM7), αmIA/IE (M5/114.15.2), αEPCAM (B40064), αCD80 (1610A1), αTCR-β (H57597), αTCR-γδ (GL3), αCD127 (SB/199), and αCCR6 (140706) mAbs (BD Biosciences; ThermoFisher; VectorLab). Intracellular staining for FOXP3 (FJK-16S) and RORγT (Q31-378) was performed after fixation/permeabilization (ThermoFisher). Stained cells were evaluated on a flow cytometer (FACSFortessa; BD Biosciences), and data analyses were performed using FACSDiva (BD Biosciences) or FlowJo software (Tree Star).

### Immunohistochemistry

Frozen thymic sections were stained, as previously described (Pouzolles et al., 2020). Briefly, sections were stained with anti–keratin 14 (1:800; AF64; Covance) and a secondary Cy3- conjugated anti-rabbit antibody (1:500; ThermoFisher) together with an Alexa Fluor 488– conjugated anti-AIRE mAb (1:200; 5H12; ThermoFisher). They were then counterstained with 49- 6-diamidino-2-phenylidole dihydrochloride (1 mg/mL), as previously reported. Images were acquired on Nikon SoRa Spinning Disk and Zeiss Axioimager Z2 microscopes and quantified using ImageJ software (National Institutes of Health).

### Digestion

Thymi were cut into approximately 1-5mm diameter pieces in RPMI at room temperature, and gently agitated using a P1000 pipet with wide-bore tips to remove thymocytes. This step was repeated until thymocytes were no longer released (supernatant remained translucent). The supernatant was then removed and thymic pieces were resuspended in RPMI (1 ml) containing DNase (0.1 mg/ml; Roche) and liberase TM (0.5 U/mg; Roche) and incubated at 37^0^C for 15 min, with the procedure repeated 3 times. Stromal fragments were agitated using wide-bore tips followed by standard P1000 tips between each incubation. After the second incubation, the supernatant was removed and diluted with PBS (5 ml) containing 2% bovine serum albumin (Sigma, ref-F7524) at RT and the remaining stromal fragments were incubated again in the enzymatic solution (1 ml). Cells were pooled after the third incubation, centrifuged at 4^0^C and filtered. Cells were then kept on ice prior to antibody staining and analysis.

### Single-Cell RNA Sequencing (scRNA-seq)

Cells were sorted into CD45.1^+^CD3^-^CD4^+^ and CD45.1^-^CD3^-^CD4^+^ population using a BD FACSAria™ sorter (BD Biosciences). The single-cell RNA sequencing was performed by the GenomiX platform (MGX) in Montpellier. Approximately 10,000 cells from each population were processed for single-cell RNA sequencing using the 10x Genomics Chromium Platform and sequenced on an Illumina NovaSeq 6000.

Sequencing data were processed using Cell Ranger (v6.0, 10x Genomics) and analyzed with NIDAP (NIH Integrated Data Analysis Platform) for clustering and differential expression.

### Statistical analyses

Statistical significance was determined using a one-way ANOVA (Tukey’s post hoc test) and unpaired t tests, as indicated (Graph Pad Software, La Jolla, CA). Data were statistically different for p≤0.05. All data are presented as means ± standard deviation (SD).

## ACKNOWLEDGMENTS

We thank all members of our lab for their scientific critique and support. The authors thank the MGX and Montpellier Ressources Imagerie facilities from Biocampus, as well as Phil Homan from the Bioinformatic platform from the NIH. The authors are grateful to Myriam Boyer and Stephanie Viala for support in cytometry experiments and to all the ZEFI staff for their support of animal experiments. This work was funded by the AFM, the ANR and INCa. A.M. was supported by a PhD fellowship from the French Ministry for reseach and Technology, C.H. was supported by a PhD fellowship from AFM and M.P. by a PhD fellowship from the LABEX EpiGenMed and the FRM. V.Z. and V.D. are supported by CNRS and C.Ha by INSERM. N.T. is supported by the NIH and was previously supported by INSERM.

