## supplemental data for "Intrathymic progenitor cell transplantation restores T cell development through ILC3-TEC crosstalk"

#### SUPPLEMENTAL MATERIAL

**Figure S1. Low engraftment of donor HSCs following intravenous administration into *Zap-70*$^{-/-}$ mice.**

**(A)** Non-conditioned *Zap-70*$^{-/-}$ mice (CD45.2$^{+}$) were transplanted by intravenous administration of WT histocompatible BM progenitors (CD45.1$^{+}$, 2x10$^{5}$) and the presence of CD45.1$^{+}$ donor cells was monitored by flow cytometry. Representative dot plots of thymi at 4, 6 and 17 weeks after transplantation are presented and the percentages of CD45.1$^{+}$ cells are indicated. Controls showing CD45.1$^{+}$ thymocytes in WT and *Zap-70*$^{-/-}$ (KO) recipients are also presented. **(B)** Quantification of the percentages of CD45.1$^{+}$ donor cells in the thymi of *Zap-70*$^{-/-}$ mice are presented at 4-17 weeks post transplantation. Each point represents data from an individual thymus. Statistical significance was determined using an unpaired 2-tailed t-test.

**Figure S2. Profiles of CD4 and CD8 lymph node cells following IT HSPC transplantation.**
(A) Non-conditioned *Zap-70*$^{-/-}$ mice (CD45.2$^{+}$) were transplanted by intrathymic administration of WT histocompatible BM progenitors (CD45.1$^{+}$, 2x10$^{5}$). Representative dot plots of CD4 and CD8 T cells in LN of WT mice and at 3, 4, 6 and 17 weeks after IT transplantation *Zap-70*$^{-/-}$ mice of are presented. (B) Quantification of CD4 and CD8 ratio in WT and IT- transplanted *Zap-70*$^{-/-}$ mice are shown at the indicated time points. Statistical significance was determined using a 1-way ANOVA with a Tukey multiple comparison test. ns, not significant

**Figure S3. Profile of the medulla following intravenous administration of HSPC into *Zap-70*$^{-/-}$ mice.**
**(A)** Thymi from WT, *Zap-70*$^{-/-}$ (KO), and IV-transplanted *Zap-70*$^{-/-}$ mice were evaluated for the presence of a thymic medulla by immunohistochemical analysis of keratin 14 (K14) and AIRE expression. Representative microscopy images of thymic tissue sections at 2, 3, 4, 6 and 17 weeks after IV injection of HSPCs are shown. **(B)** Quantification of the numbers of large K14$^{+}$ zones (>10 000/ mm$^{2}$; left panel) and the number of AIRE$^{+}$ cells per mm$^{2}$ (right panel) are presented for the indicated conditions. Quantifications were performed in medulla from at least 3 individual mice. **(C)** The presence of CD80$^{hi}$MHCII$^{hi}$ mTEC was evaluated by flow cytometry following digestion of thymi from WT, *Zap-70*$^{-/-}$, and IV-transplanted *Zap-70*$^{-/-}$ mice, at the indicated time points. mTEC were evaluated on gated EPCAM$^{+}$ UEA-1$^{+}$ cells and representative plots are shown. **(D)** Quantification of CD80$^{h}$iMHCII$^{hi}$ mTEC in WT and *Zap-70*$^{-/-}$ mice at the indicated time points after IV HSPC transplantation (n=4-8 mice per group). Means ± SD are presented. Statistical significance was determined using a 1-way ANOVA with a Tukey multiple comparison test.

**Figure S4. scRNAseq analysis of immature recipient and donor CD3$^{-}$CD8$^{-}$ thymocytes.**
**(A)** Spectral tSNE plots of all scRNAseq-analyzed CD3$^{-}$CD8$^{-}$ thymocytes annotated by cell-type identity. **(B)** Proportions of immune cells presents in all cells. The percentages of DN1, DN2, DN3, DN4, ILC3, and LTi thymocytes are presented. **(C)** Spectral tSNE plots of FACS-sorted CD3$^{-}$CD8$^{-}$ thymocytes colored by the provenance of the cells (recipient vs donor, right) and by clusters (right). **(D)** Percentages of FACS-sorted CD3$^{-}$CD8$^{-}$ donor and recipient thymocytes that are in each cluster (left) and the percentages of cells within each cluster that are from the recipient as compared to the donor (right).

### Figure S1

A

B

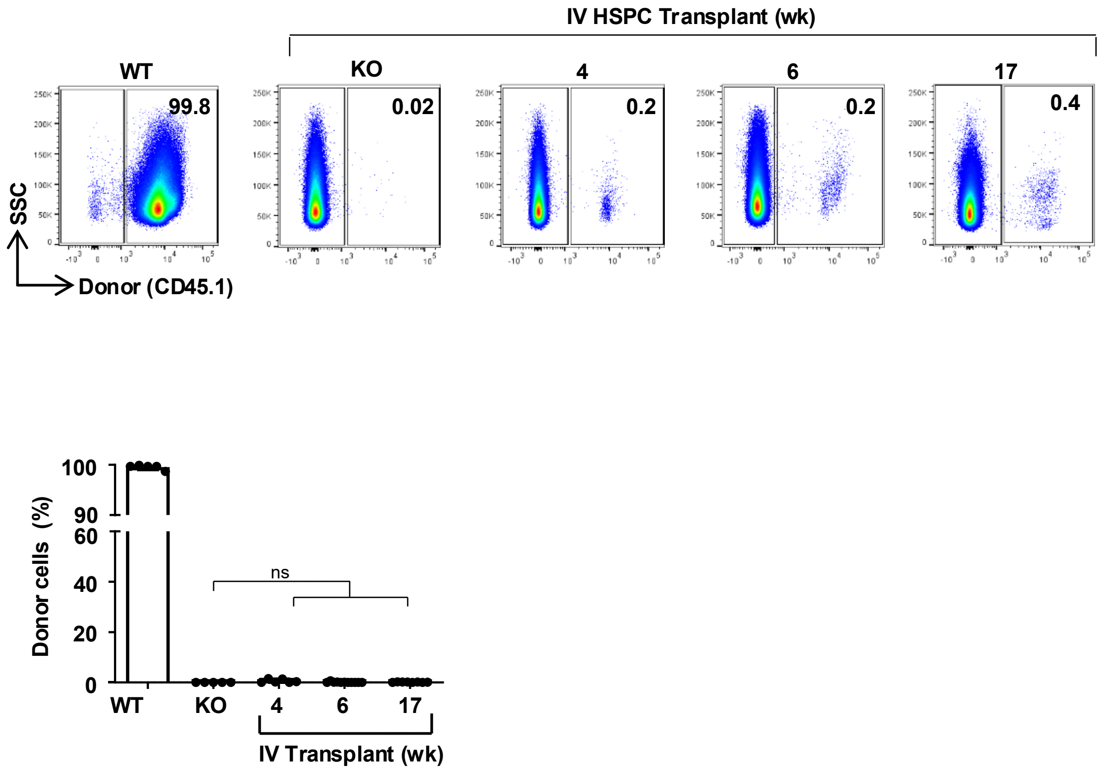

IV HSPC Transplant (wk)
WT
KO
4
6
17
99.8
0.02
0.2
0.2
0.4
SSC
Donor (CD45.1)
Donor cells (%)
100
90
60
40
20
0
ns
WT
KO
4
6
17
IV Transplant (wk)

### Figure S2

## A

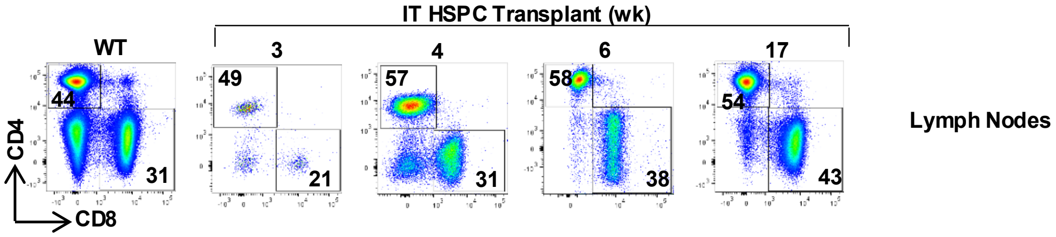

IT HSPC Transplant (wk)
WT
3
4
6
17
44
31
49
21
57
31
58
38
54
43
CD4
CD8
Lymph Nodes


## B

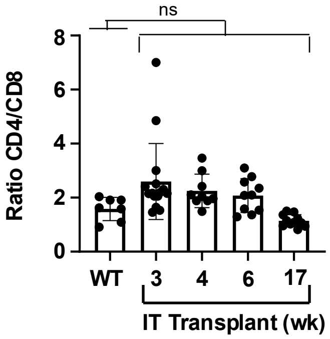

ns
Ratio CD4/CD8
8
6
4
2
0
WT
3
4
6
17
IT Transplant (wk)

### Figure S3

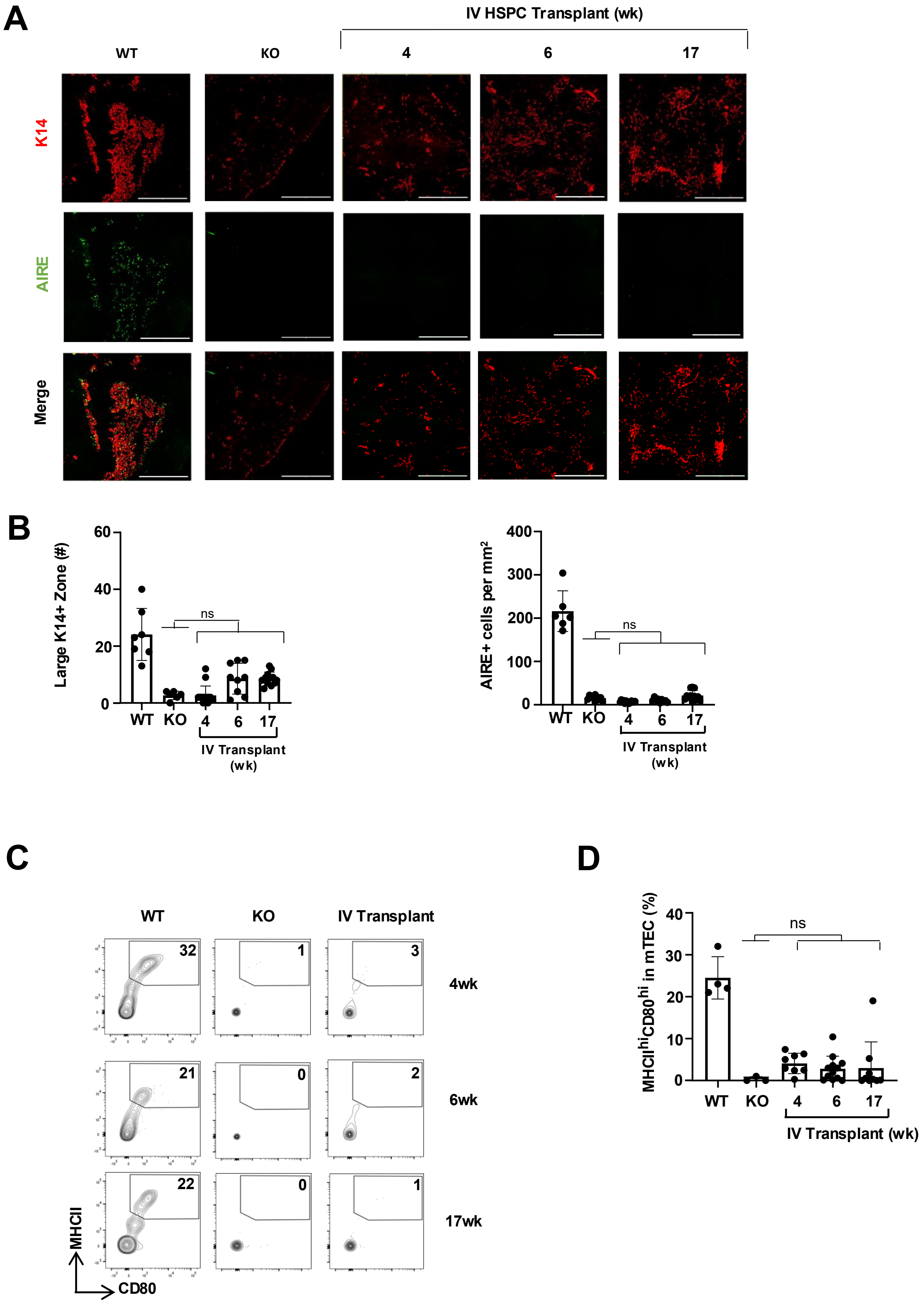
A
IV HSPC Transplant (wk)
WT
KO
4
6
17
K14
AIRE
Merge
B
Large K14+ Zone (#)
60
40
20
0
ns
WT KO 4 6 17
IV Transplant (wk)
AIRE+ cells per mm²
400
300
200
100
0
ns
WT KO 4 6 17
IV Transplant (wk)
C
WT
KO
IV Transplant
32
1
3
4wk
21
0
2
6wk
22
0
1
17wk
MHCII
CD80
D
MHCII^hi CD80^hi in mTEC (%)
40
30
20
10
0
ns
WT KO 4 6 17
IV Transplant (wk)

### Figure S4

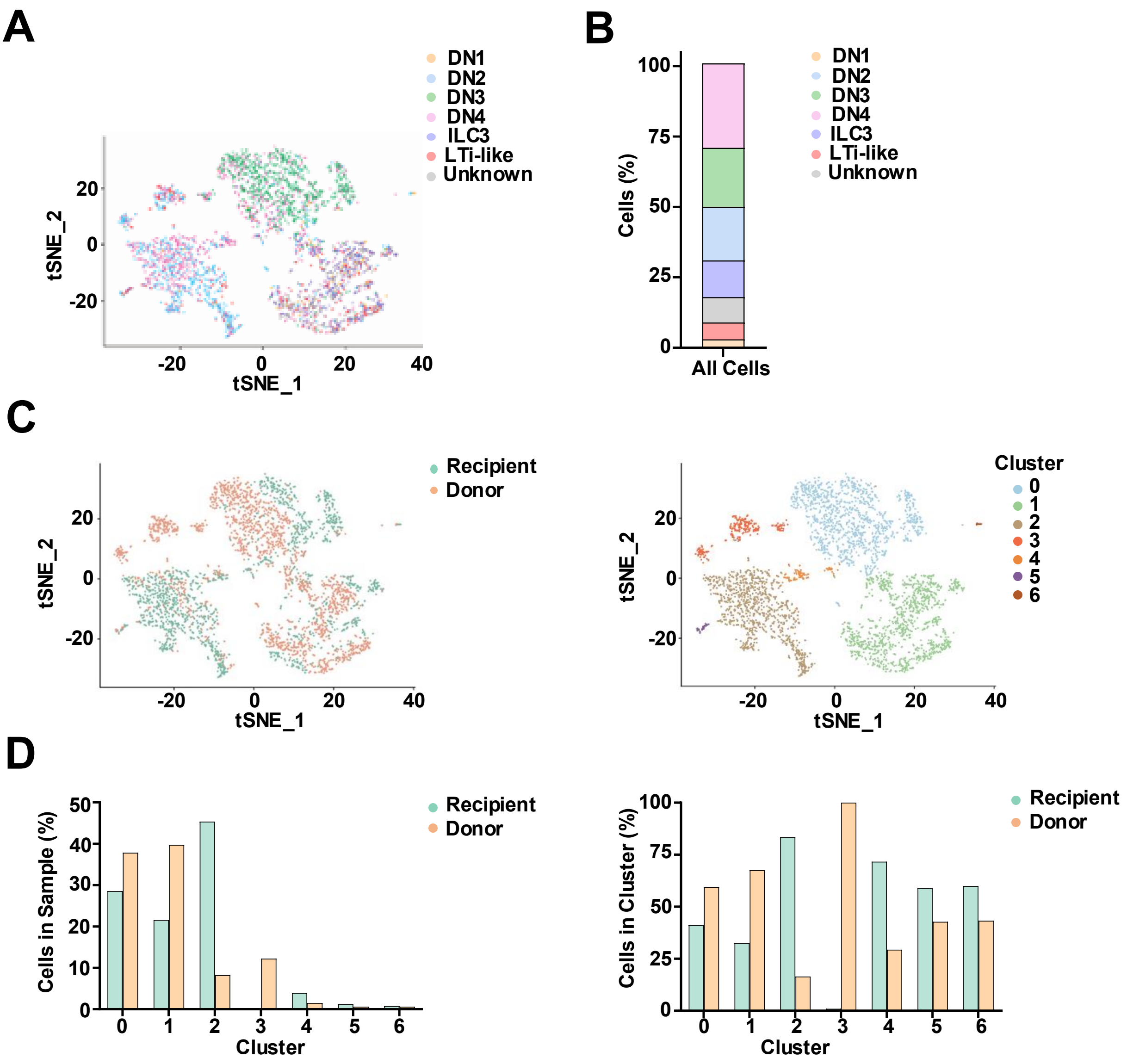
A
DN1
DN2
DN3
DN4
ILC3
LTi-like
Unknown
tSNE_2
tSNE_1
20
0
-20
40
B
Cells (%)
100
75
50
25
All Cells
C
Recipient
Donor
Cluster
0
1
2
3
4
5
6
D
Cells in Sample (%)
Cells in Cluster (%)
Cluster

### Figure S5

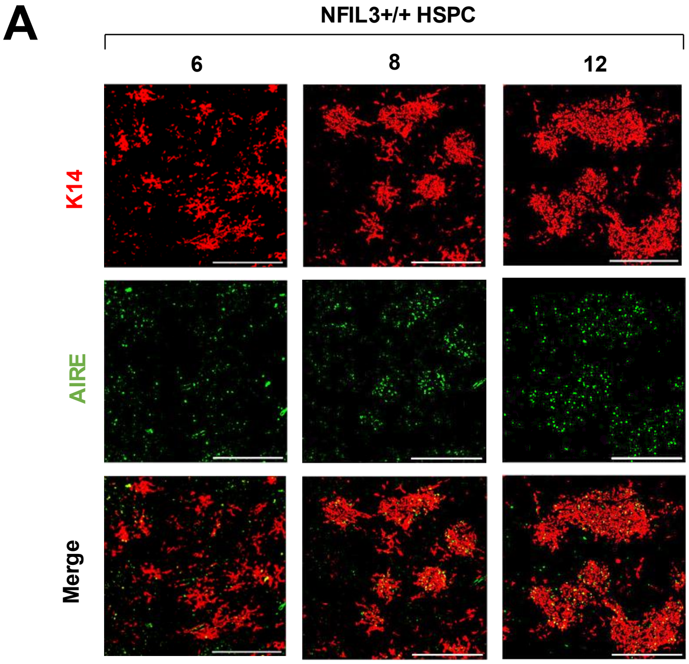
